# Host-microbiome coevolution promotes cooperation in a rock-paper-scissor dynamic

**DOI:** 10.1101/689299

**Authors:** Ohad Lewin-Epstein, Lilach Hadany

## Abstract

Cooperation is a fundamental behavior observed in all forms of life. The evolution of cooperation has been widely studied, but almost all theories focused on the cooperating individual and its genes. We suggest a different approach, taking into account the microbes carried by the interacting individuals. Accumulating evidence reveal that microbes can affect their host wellbeing and behavior, yet hosts can evolve mechanisms to resist the manipulations of their microbes. We thus propose that coevolution of microbes with their hosts may favor microbes that induce their host to cooperate. Using computational modeling, we show that microbe-induced cooperation can evolve and be maintained in a wide range of conditions, including when facing hosts’ resistance to the microbial effect. We find that host-microbe coevolution leads the population to a rock-paper-scissors dynamic, that enables maintenance of cooperation in a polymorphic state. This theory may help explain occurrences of cooperation in a wide variety of organisms, including in cases that are difficult to explain by current theories. In addition, this study provides a new perspective on the coevolution of hosts and their microbiome, emphasizing the potential role of microbes in shaping their host behavior.

## Introduction

Cooperative behavior, such that confers a fitness cost to the acting individual, while providing a benefit to its partner, is a fundamental behavior, present in all levels of organization – from bacteria to communities of multicellulars. As such, the evolution of cooperation by the means of natural selection presents a puzzle [1-7].

A major class of models was introduced by Hamilton [8], who suggested that natural selection may favor a gene that induces cooperative behavior, if directed towards kin, which are likely to carry other copies of the same gene [9-16]. Another class of explanations suggested reciprocity as an explanation for cooperation [17], focusing on the benefit to the cooperating individual. According to this theory, when interactions are recurring, conditional cooperative behavior can evolve, if directed towards individuals that cooperated in previous rounds – even if they are not relatives [2, 18-22]. This includes direct reciprocity (A helps B and B helps A), and indirect reciprocity (A helps B and C helps A, based on the reputation of A) [23, 24]. Cooperation was further suggested to be favored as a signal for the quality of the cooperating individual, which is eventually rewarded with increased social status, mating success, etc. [25, 26]. More recent work investigated the effect of population structure and viscosity – affecting, among other things, the probability of repeating interactions and interactions among kin – on the evolution of cooperation [27-31].

The vast majority of theories offering an explanation for the evolution of cooperation share a common attribute, focusing on the genes of the cooperating individual, namely on traits that affect the tendency to cooperate and are transmitted from parent to offspring. Recently, we suggested an alternative explanation, focusing on the *microbes* carried by the interacting individual, that can be transmitted both from parent to offspring and between interacting hosts. We showed that microbes that induce host cooperation can evolve under wide conditions [32]. Here we combine the two approaches and consider both the host genes and its microbiome coevolving with respect to cooperation.

Almost all organisms carry microbes, that can have dramatic effects over their host wellbeing and behavior [33-39]. Recent evidence shows that the gut microbiome can affect the brain via the microbiome-gut-brain-axis [40-42], potentially affecting brain development, cognitive function, and behaviors such as social interactions and stress [43-48]. In light of this evidence, we raised the hypothesis that microbes can also affect the tendency of their hosts to cooperate.

In cases where microbes perform a manipulation on their host, a conflict of interest may arise. This could lead to evolution of the hosts by acquiring traits that negate the microbial manipulation and provide resistance [49-52]. In many cases, such a resistance may itself confer a cost to the host [53-56], generating complex dynamics of coevolution of the hosts and their microbes.

In addition to the ability to affect host behavior, microbes have varied dispersal abilities between hosts. Similarly to genes, microbes can be transmitted vertically, namely inherited, from one individual to its offspring [57-61]. Yet, as opposed to genes of multicellular organisms, microbes can also be transmitted horizontally, between interacting individuals [62-65]. Due to the ability of microbes to transmit horizontally, they can benefit from inducing their host to help another host, that could be inhabited by their transmitted kin – even when the hosts are not related. In that respect, our theory corresponds to the theory of kin selection, where the relevant “kin” are not the interacting individuals, but rather the microorganisms that inhabit them.

Here we study host-microbe coevolution, and analyze the conditions that allow the evolution of microbe-induced cooperation in a population of hosts that can evolve resistance to the microbial effect. In our framework we consider the abilities of the microbes to be transmitted both vertically and horizontally, as well as the cost and benefit of cooperation, and the cost of host resistance to the microbial effect. We find that under a wide range of parameters, microbe-induced cooperation facing host resistance generates a rock-paper-scissor dynamic in the population. Thus, polymorphism with respect to host behavior is preserved, allowing the evolution and maintenance of cooperation, and supporting the theory of microbe-induced cooperation.

## Results

### Model description

We model a population of asexual haploid hosts, each carrying one type of microbes, *α* or *β*. Microbes of type *α* increase the tendency of their hosts to cooperate, while microbes of type *β* don’t have any effect over their host behavior. In addition, a bi-allelic locus at the host genome determines the susceptibility of the host to the microbial effect. Hosts carrying allele *S* are susceptible to the microbe’s effect, and act cooperatively when carrying microbe *α*. Hosts carrying allele *R* are resistant to the microbial effect and do not act cooperatively regardless of the microbe they carry. This resistance confers a fitness cost of 0 < *δ* < 1 (we investigated also the case where the resistance cost depends on the type of microbe type the host carries, and found qualitatively similar results. See the Methods section as well as SI1-3). We model a haploid population with four different types of hosts: *αS, αR, βS, βR*, defined by the combination of microbes (*α*/*β*) that the host carries, and its allele (*R*/*S*).

Each generation, hosts pair randomly and interact, with a prisoner’s dilemma payoff (see Fig. 1a). During interaction the fitness of the interacting hosts changes according to their behavior: an individual that behaves cooperatively (type *αS* only) pays a fitness cost 0 < *c* < 1 while its partner receives a benefit *b* > *c*. A selfish individual doesn’t pay the fitness cost, but receives the benefit if its partner is a cooperator. In addition, horizontal transmission of the microbes might occur during interaction (see Fig. 1b). We denote by *T*_*α*_ the probability of microbes of type *α* being transmitted from one host to the other during interaction, establishing and replacing the resident microbes, and similarly with *T*_*β*_. We assume that transmission of one microbe is independent of the other microbe, and when both occur, they occur simultaneously. We note that *T*_*α*_ and *T*_*β*_ encompass the probability of completing the entire transmission process: traversing the physical barrier, competing with the native microbial community, and establishing a colony.

**Figure 1.**
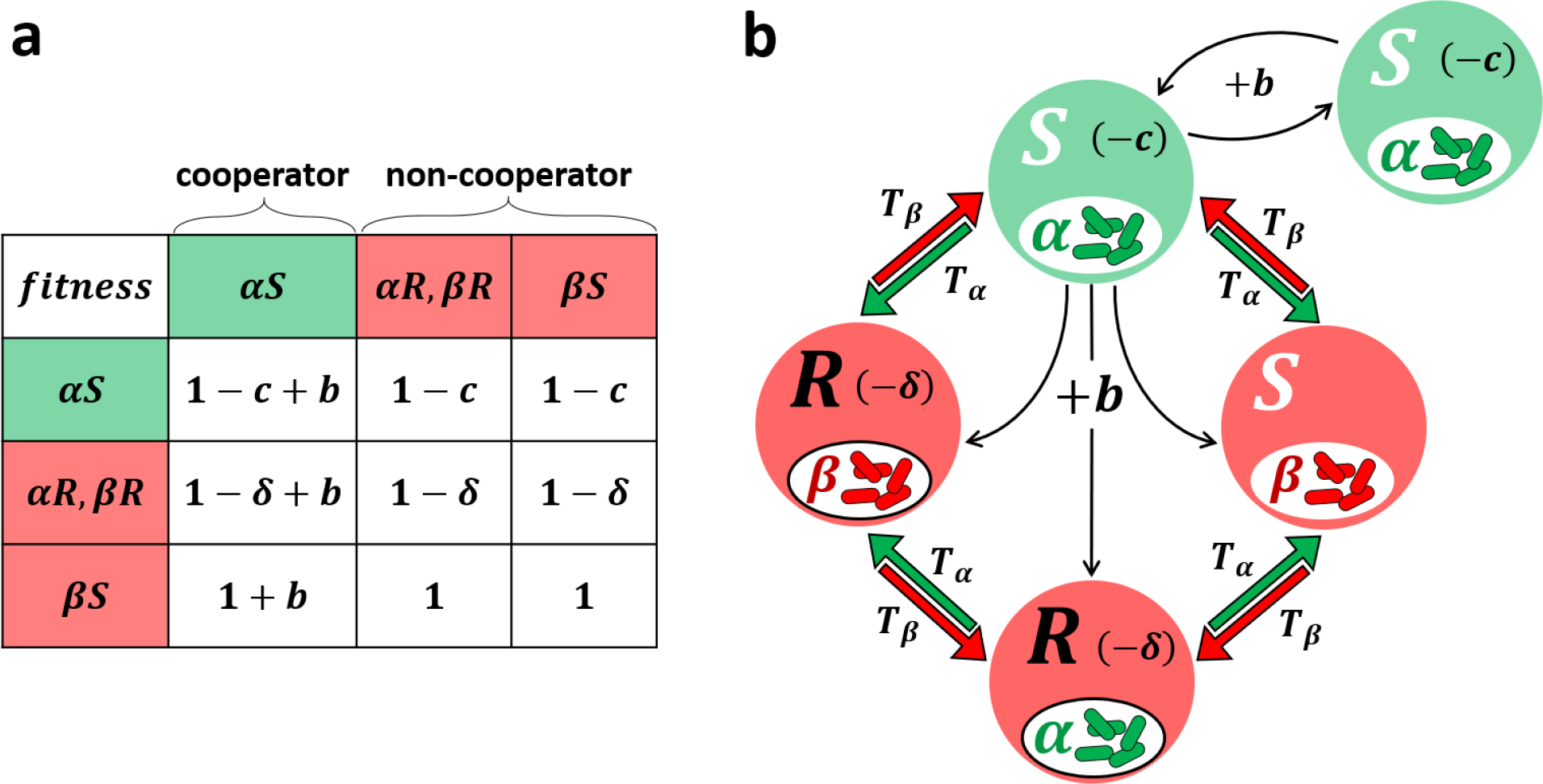
Model illustration. Each individual hosts one type of microbes, either *α* (inducing cooperation) or *β* (no effect). In addition, each host carries either allele *S* (susceptible to microbial effect) or *R* (resistant). Thus, only *αS* individuals are cooperators. Carrying allele *R* also confers a fitness cost of *δ*. When two individuals interact, their fitness can change as a result of the interaction. α*S* individuals cooperate: in each interaction they pay a fitness cost of 0 < *c* < 1, and their partners receive a fitness benefit *b* > *c*. Hosts of all other types don’t cooperate. In addition, horizontal transmission of microbes may occur during interaction, regardless of the alleles that the hosts possess. We mark by *T*_*α*_ the probability of microbes of type *α* being transmitted to the other host, establishing and replacing the resident microbes, and similarly with *T*_*β*_. **(a)** Fitness matrix showing the fitness of each host, according to allele, microbe, and interaction partner**. (b)** Possible interactions that yield fitness change, microbe transmission, or both. In brackets are the fitness costs for the individuals: −*δ* for hosts with allele *R*, and −*c* for cooperators (*αS* hosts). Black arrows represent the fitness benefit (*b*) that cooperators provide to their partners. Colored arrows (red and green) represent the probability for microbial horizontal transmission during interactions.

We use discrete models, with non-overlapping generations. At the end of each generation, after interactions and horizontal transmissions take place, the hosts reproduce according to their fitness. Both genes and microbes are vertically transmitted from hosts to their offspring, and the offspring host generation replaces the parent generation. We investigate this general model using two approaches. First, in *Deterministic model* section, we analyze the dynamics of a fully-mixed infinite population. Second, in *Stochastic models* section, we use computational simulations to analyze the dynamics of a finite population, both fully-mixed and spatially structured, and account for stochastic effects and mutations. The additional details of each model are included below in the relevant sections and in the Methods.

### Deterministic model

We first consider an infinite, fully-mixed population, where each individual has an equal probability to interact with any other individual in the population. Every generation the population is randomly divided into pairs and each pair interacts once. During these interactions, the fitness of the hosts is determined by the payoff of the interaction, and the microbes can be transmitted between the interacting hosts (see Fig. 1). Cooperating individuals are only those that carry both allele *S* and microbe *α*. We describe the change in the frequencies of the different host types from one generation to the next using four iterative equations (see equations 4-7 in the Methods). By analyzing this system of equations, we study the conditions that allow the evolution and maintenance of cooperative behavior.

We first find that cooperation can evolve (i.e., *αS* gene-microbe combination can increase from rarity) only when:

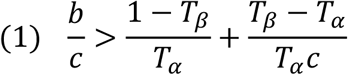

see Figures 2a, b, and SI2-3. This result is consistent with [32], where a similar condition determines the evolution of a microbe inducing cooperation in a population of hosts that are all susceptible to the microbe’s effect. Intuitively, the conditions allowing the evolution of cooperation in the presence of host resistance (here) are never wider than the conditions in a susceptible population [32]. We thus continue by assuming (1) is satisfied and analyzing the additional conditions that allow host susceptibility to the microbes (allele *S*) to increase in frequency. Denoting the proportions of hosts carrying allele *S* (*αS* and *βS* hosts), and of hosts carrying both allele S and microbe *α* (*αS* hosts) by *x*_*S*_ and *x*_*αS*_, respectively, we find that the proportion of allele *S* increases from one generation to the next if and only if (see analysis in SI3):

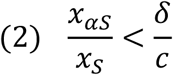

We first see that if *δ* > *c*, condition (2) is always satisfied and thus the proportion of allele *S* increases to fixation for any *x*_*S*_ > 0. Quite expectedly, we see that when the cost of cooperation (*c*) is smaller than the cost of host resistance to the microbial effect (*δ*), cooperation will fixate in the population (area *II* in figure 2, panels a and b). Furthermore, we find that when *δ* < *c*, cooperation can evolve and be maintained in polymorphism, and that a polymorphic equilibrium, when it exists, maintains 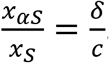.

**Figure 2.**
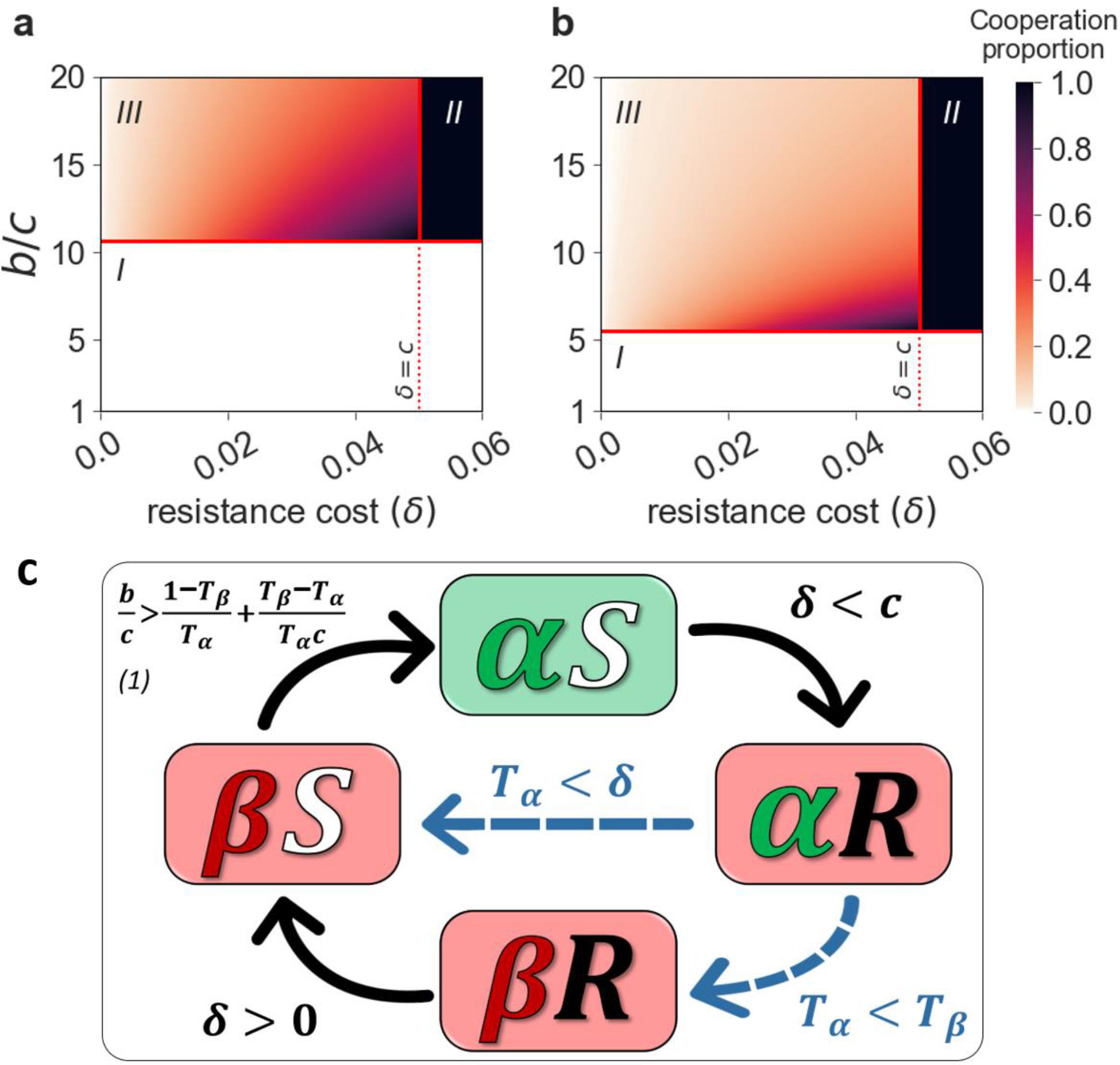
Cooperation can be maintained at intermediate levels even in the presence of host resistance to the microbial effect. **(a**,**b)** We plot (color coded) the expected proportion of cooperative hosts (*αS*), as a function of the *b*/*c* (y-axis) and of *δ* (x-axis) for *c* = 0.05, *T*_*β*_ = 0.25, *T*_*α*_ = 0.75 · *T*_*β*_ **(a)**, *c* = 0.05, *T*_*β*_ = 0.25, *T*_*α*_ = 0.9 · *T*_*β*_ **(b)**. Cooperation goes extinct when below the horizontal dashed line representing condition ***(1)*** (area ***I***, white). Above that threshold, cooperation can either go to fixation (when *δ* > *c*, area ***II***, black), or be maintained at intermediate levels (when *δ* < *c*, area ***III***). In the latter case, the proportion of cooperators either converges to the equilibrium or oscillates around it. **(c) Rock-paper-scissors game of cooperation.** Based on invasion analysis (see SI2), the figure illustrates conditions that allow the invasion of a rare type to a population dominated by another type. When *δ* < *c, αR* hosts can invade an *αS* population, due to the lower cost of resistance in comparison with the cost of cooperation; an *αR* population can be invaded by either *βR* individuals (if *T*_*β*_ > *T*_*α*_) or *βS* individuals (if *δ* > *T*_*α*_); as long as *δ* > 0 then *βS* individuals invade a *βR* population, as they don’t pay the cost of resistance; finally, if 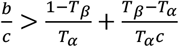 (condition ***(1)***) then *αS* individuals can invade a *βS* population. Note that there are two possible cycles that allow the maintenance of cooperation: *αS* → *αR* → *βS* → *αS* and *αS* → *αR* → *βR* → *βS* → *αS*. Altogether, cooperation will be maintained as long as all the conditions represented by black arrows are satisfied, in addition to at least one of the conditions represented by dashed blue arrows.

When *δ* < *c, αS* hosts bear an inherent disadvantage – they pay a fitness cost of *c*, while the rest of the hosts pay lower costs of either *δ* (*αR* and *βR* hosts), or no cost at all (*βS* hosts). Yet, we find that polymorphic equilibria exist and cooperation can evolve under a wide parameter range and be maintained at intermediate levels (area *III* in figure 2, panels a and b). The proportion of cooperators at this polymorphic equilibrium increases with the cost of resistance (*δ*), but is bounded: cooperators can reach up to *δ*/*c* from the proportion of hosts carrying allele *S* in the population. Counter intuitively, above the *b*/*c* threshold (of condition (1)) the proportion of cooperators at equilibrium decreases with *b*/*c*, due to the evolution of resistance.

To identify the conditions that allow polymorphism, we investigated the stability of the four trivial equilibria (namely, the fixations of the four host types). Invasion analysis (detailed in SI2) revealed that when 0 < *δ* < *c*, cooperation evolves whenever:

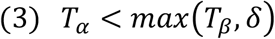

When (1) and (3) are satisfied and 0 < *δ* < *c*, no equilibrium that involves extinction of some of the host types, is stable. Under these conditions, cooperation is maintained at rock-paper-scissor dynamics (see Figures 2c and 3). Condition (3) is somewhat counter-intuitive: for example, when *α* microbes have a transmission disadvantage (*T*_*α*_ < *T*_*β*_), it can facilitate the evolution of cooperation by hindering the evolution of resistance.

**Figure 3.**
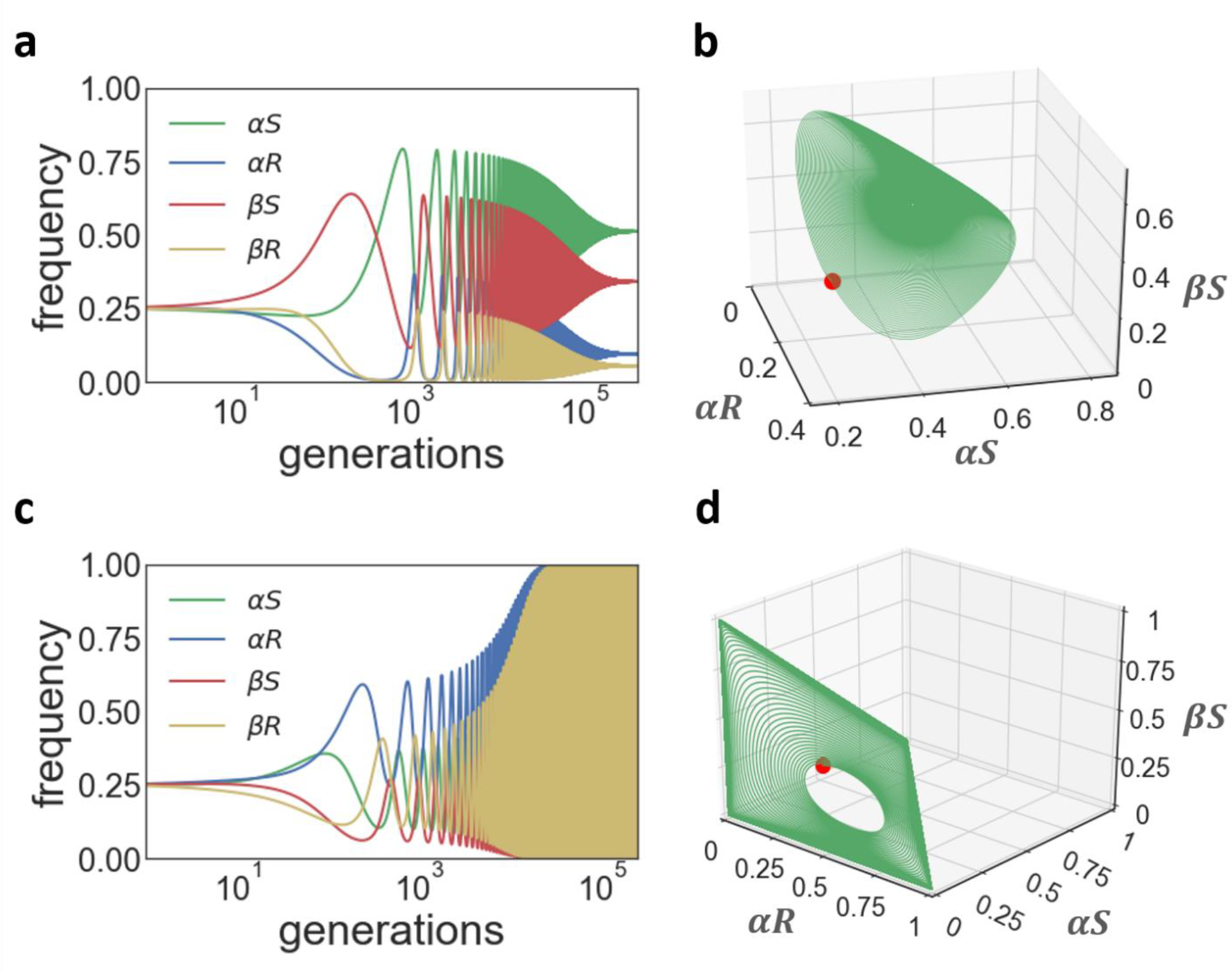
Oscillations of cooperation can either converge or diverge. We plot the frequencies of the four individual types in the population with time (panels **a, c**) and in a 3D plane (panels **b, d**) for *c* = 0.05, *δ* = 0.03, *T*_*β*_ = 0.25, *T*_*α*_ = 0.9*T*_*β*_ (based on iterations of equations (4-7), see Methods). **(a**,**b) *b***/***c*** = **6. Converging oscillations**. The red dot in **(b)** represents the initial state of the population, and the population spirals towards the focal point until it fixates. **(c**,**d) *b***/***c*** = **10. Diverging oscillations.** The red dot in **(d)** represents the initial state of the population, and the frequencies oscillate and slowly diverge. In all panels the initial state of the population includes all four host types at equal proportions.

Interestingly, we found that the behavior of the polymorphic system is oscillatory, and the population can either converge with oscillations towards the equilibrium (Figure 3, panels a and b) or oscillate chaotically around the equilibrium (Figure 3, panels c and d), depending on the different parameters and on the initial conditions of the population. In a population undergoing such chaotic oscillations, cases of near-fixation of one of the types are frequent. The behavior of a finite population undergoing similar dynamics is thus intriguing: could cooperation still be maintained?

### Stochastic models

So far, we considered an infinite population where the dynamics are deterministic. Here we study the evolution of microbe-induced cooperation in finite populations, subject to stochastic effects. We constructed a stochastic simulation modeling a population of 10,000 interacting hosts (see simulation workflow in the Methods section). Similarly to the analytic model, each host carries microbes of type *α* or *β* and allele *R* or *S*, and horizontal transmission of the microbes can occur during interactions. Each generation individuals are paired randomly, and one interaction takes place between the two members of each pair. We start with a fully mixed and mutation free population, corresponding to the analytical model.

The simulation results strongly agree with the deterministic fully-mixed model with regard to the conditions that lead to the fixation or extinction of cooperation, but there is a difference with regard to the conditions that allow polymorphism. While in the deterministic model we found long term polymorphism, in the finite population cooperation goes extinct in many of the parameter sets (compare area ***III*** of Figures 2a and 2b to area ***III*** in Figure 4a).

**Figure 4.**
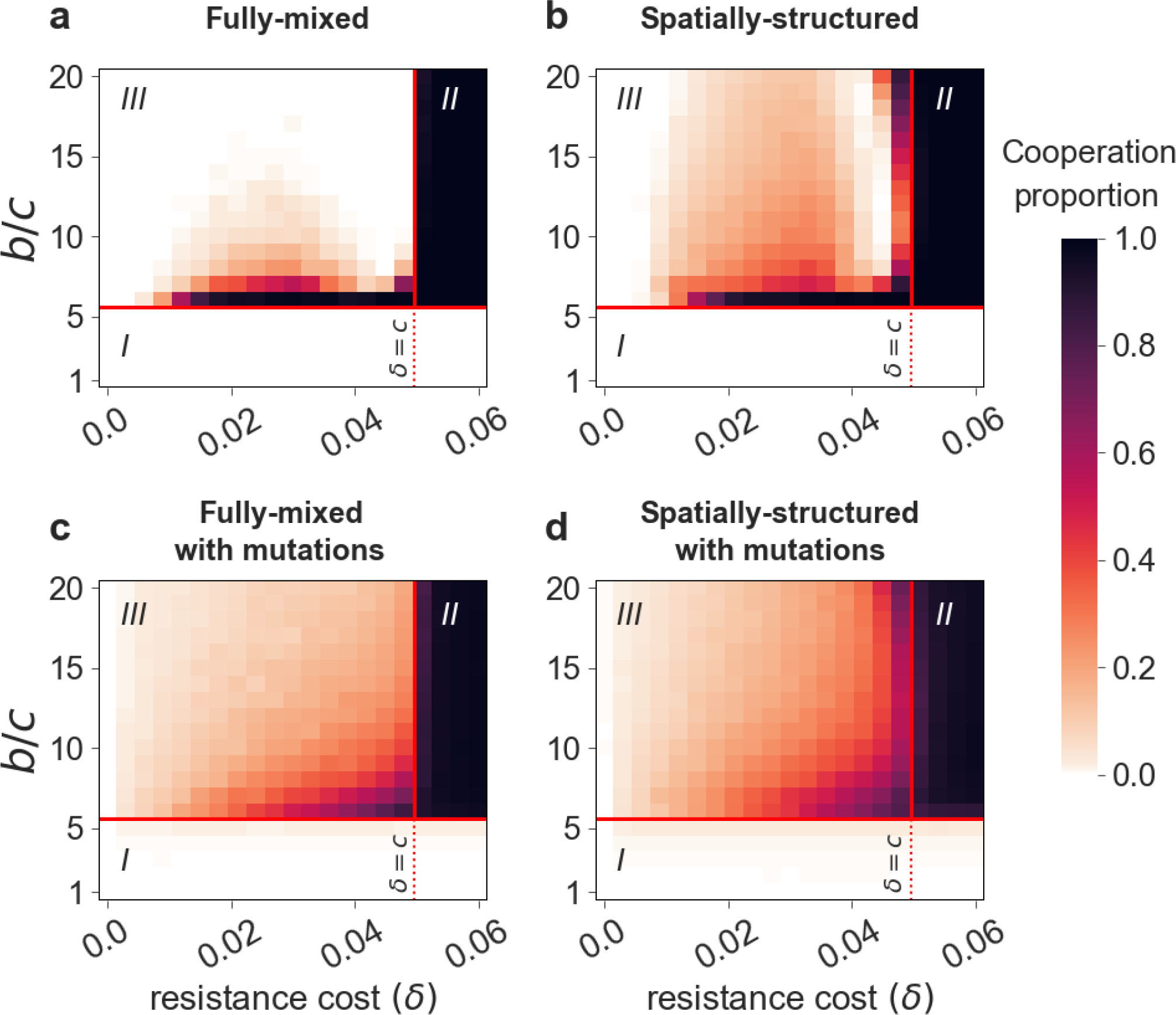
Mutations and spatial structure help maintaining cooperation in the face of host resistance and finite populations. The proportion of cooperators after 5,000 generations is plotted as a function of the *b*/*c* ratio on the y-axis and *δ* on the x-axis. The color of each site represents the average of 200 stochastic simulation runs. Panels **(a)** and **(c)** show the results of fully-mixed populations while panels **(b)** and **(d)** show the results of spatially-structured populations. Panels **(a)** and **(b)** show the dynamics without mutations while panels **(c)** and **(d)** show results of populations with mutation rates of *µ* = 10^−4^ in all directions (*α* ↔ *β and S* ↔ *R*). Consistent with the analytic results, we find that cooperation goes extinct if (1) is not maintained, and cooperation fixates if (1) is maintained and *δ* > *c*. We find that both mutations and spatial structure significantly widen the parameter range allowing the maintenance of cooperation in polymorphism. Simulation parameters: *T*_*β*_ = 0.25, *T*_*α*_ = 0.9*T*_*β*_, *c* = 0.05. The results were robust to different mutation rates of *µ* = 10^−3^ and *µ* = 10^−5^ and to a different time limit of 10,000 generations (see SI4).

There are several mechanisms that can maintain polymorphism in an oscillating population, such as mutations, spatial structure, and others [66-70]. Mutations keep generating hosts and microbes of all types in the population, and by that rescue rare or extinct types, giving them additional chances to spread. Spatial structure on the other hand limits both dispersal and interaction to the local scale, thus decreasing the strength of competition between the different host types – even when a type is common in the population, it is not common in every patch. As a result, the strength of oscillations decreases and all host types are maintained for a long time.

We therefore extended the simulation to account for both mutations and spatial structure (see details in Methods section). Mutations were modeled as a random change (with rate *µ*) in an offspring allele or/and microbe type, relative to its parent. Spatial structure was modeled similarly to [71], using a 2D-lattice of size 100 × 100, where each site is inhabited by one host. Differently from the fully-mixed model, in the spatially structured population the interactions occur only between individuals of neighboring sites, and selection is local as well. After the hosts interact, reproduction takes place, and each site is inhabited by an offspring of a parent from the neighborhood (3X3 sites, or less if adjacent to the border), chosen according to the parent’s fitness. The offspring carries the same allele and microbes’ type as its parent, to the point of mutations.

We find that both mutation and spatial structure dramatically widen the range of parameters that allow the maintenance of cooperation in polymorphism, when *δ* < *c* and (1) is satisfied (compare area ***III*** of Figures 4b and 4c to 4a; see also supplementary movie). The combination of both spatial structure and mutations allows even higher proportions of cooperation (Figure 4d). All the stochastic simulations (with or without mutation and spatial structure) were also consistent with the deterministic results regarding the parameter range where cooperation goes extinct or fixates.

## Discussion

Our results demonstrate that cooperation induced by microbes can evolve in a host population under a wide range of parameters including the case where the hosts coevolve and acquire resistance to the microbial induction of cooperation. Although cooperative hosts bear an inherent disadvantage, the host-microbe coevolution generates a rock-paper-scissor dynamic in the population, that enables the evolution and maintenance of cooperation. In addition, we find that the cost of resistance to the microbial effect (*δ*) is a crucial factor. If this cost is higher than the cost of cooperation (*c*) then cooperation can fixate in the host population, and if not, cooperation can be maintained at intermediate proportion. We also find that in finite-size populations, the oscillatory dynamic, generated by the rock-paper-scissor game, leads in many cases to extinction of cooperation. Consistent with other theoretical and experimental results, we show that spatial structure of the population, and mutations of both the microbes and the hosts, enables long-term maintenance of polymorphism. Our model suggests an explanation for the evolution and maintenance of cooperative behavior in a wide variety of organisms, and can shed light over cases of long term polymorphism with respect to social behavior.

Our framework can be extended in several directions. We currently model a binary microbe (either *α*, or *β*) and host alleles (either *S* or *R*). In natural populations, hosts’ behavior might be affected by the composition of the microbial community and by several loci in the genome. In addition, the host behavior can be modeled as a continuous trait, where the level of cooperation is varied, or condition-dependent. Interesting host-microbes dynamics can arise when the cooperative behavior is applied only under certain circumstances, e.g. when the hosts are under stress, similarly to other stress-induced behaviors [72-75]. It would also be interesting to examine the evolution of the rates of microbe horizontal and vertical transmission, in the context of microbe-induced cooperation. Once cooperation is common in the population, it can further affect the future evolution of the population [76, 77].

Note that in this work we define ‘microbe’ in the most general sense – an element that can affect host behavior, and can be transmitted both horizontally and vertically. Thus, our results are relevant to any element that applies to this category (e.g. plasmids, viruses, macro-parasites, epi-genetic elements etc.). There is already some evidence suggesting that microbes can affect cooperative behavior in their host. It was shown that bacterial genes responsible for public good secretion are prevalent on mobile elements that can be both horizontally and vertically transmitted between host cells [78, 79]. Among eukaryotic hosts, lactobacillus is a promising candidate: accumulating evidence demonstrates that lactobacillus decreases stress-induced anxiety-like behavior, which can increase the tendency of its host to interact with other individuals [80-82], and may further promote cooperative behavior [83, 84]. In plants, carbon sharing among trees [85, 86] is, at least partially, mediated by mycorrhizal fungi that form networks connecting neighboring tree roots. This behavior can be considered as cooperation among trees, induced by their fungi. Our results suggest that such effects may be common in many other systems as well.

This work can also be viewed in a somewhat different context of gene-culture coevolution [87, 88]. Similarly to microbes, culture also affects behavior, and is transmitted both vertically and horizontally. Moreover, like microbes, culture interacts and coevolves with the genome [89, 90]. In such a scenario, individuals cooperate or not based on the set of practices and beliefs they possess. Parents teach their offspring these cultural-behavioral traits, but they can also be transmitted horizontally, namely, an individual might imitate the behavior of another individual, especially its interaction partner [91]. “Resistance” to culture can be a relevant and important component. Similarly to the case of host alleles that affect the susceptibility to microbes, genes can also affect the tendency of individuals to follow cultural rules.

This study suggest that microbes may have a significant role in shaping their hosts behavior. It also demonstrates how a conflict between hosts and their microbes, portrayed by the ability of the hosts to evolve resistance to the microbial effect, can lead the population to a rock-paper-scissor game. This rock-paper-scissor game enables long-term maintenance of cooperation, at intermediate levels. These results strengthen the theory of microbe-induced cooperation. Furthermore, this model may help explain occurrences of cooperation that are difficult to explain by current theories, and it predicts that altering the composition of microbial communities (e.g. by antibiotics [92-94], probiotics [95, 96], pesticides [97] and herbicides [98], etc.) may affect the hosts’ social behavior. This work provides verifiable predictions that can be tested in future experimental efforts.

## Methods

### The host-microbe co-evolution model

In this section we describe the full model used for this study, and present the core equations. We model an asexual population of hosts, and assume that each host carries one of two microbe types. Microbes of type *α* increase the tendency of their hosts to cooperate, while microbes of type *β* don’t affect the host behavior. In addition, we model a host gene that determines the susceptibility of the host to the microbial effect. Hosts carrying allele *S* are susceptible to the microbe’s effect and act cooperatively when carrying microbe *α*. Hosts carrying allele *R* are resistant to the microbial effect and act selfishly at all times, but this resistance confers a fitness cost. In this section we consider a more general modeling of the resistance cost, by allowing two different resistance costs, that depend on the type of the carried microbe. We denote by *δ*_*α*_, *δ*_*β*_ > 0 the fitness cost of resistance, for hosts carrying microbes of type *α* and hosts carrying microbes of type *β*, respectively. We thus have in our model a haploid population with four different types of hosts: *αS, αR, βS, βR*, defined by the combination of alleles (*R*/*S*) and microbes (*α*/*β*). We assume that the cost of resistance is applied before any horizontal transfer occurs. We model the interactions payoff and horizontal transmissions as defined in the **Results** section.

We mark by *x*_*αS*_, *x*_*αR*_, *x*_*βS*_, *x*_*βR*_ the proportions of the host types in the current generation, and calculate 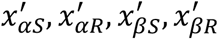, the proportions of the host types in the next generation:

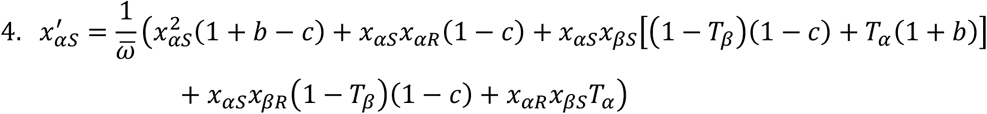

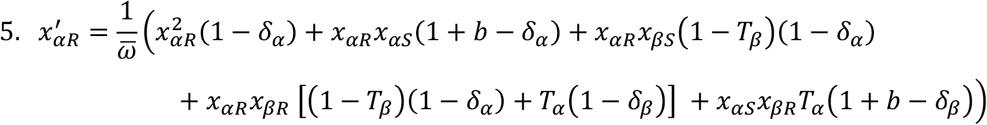

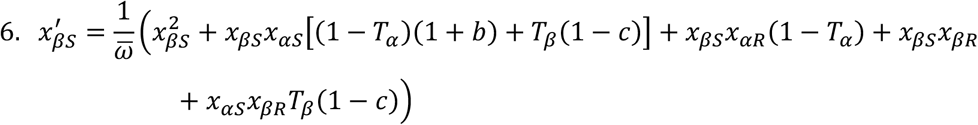

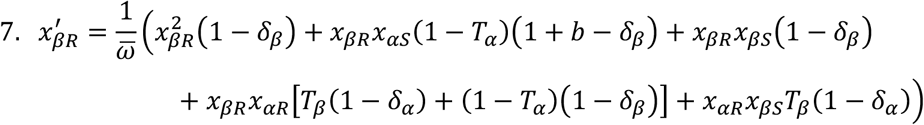

where

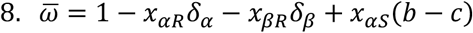

Further analysis of these equations, the equilibria and the stability analysis are presented in the Supplementary Information.

### Stochastic Simulation Work Flow

#### Fully-mixed population

We programmed an agent-based simulation, where we follow *N* = 10,000 hosts. Each host is defined by the allele (*S*/*R*) and microbe (*α*/*β*) it carries. Each generation, the hosts are divided to interacting couples. During the interaction, hosts that carry microbe *α* and allele *S* cooperate: they pay a fitness cost, *c*, and their partner receives a fitness benefit, *b*, regardless of its microbe-allele combination. Hosts with microbe-allele combination other than *αS*, don’t cooperate, and thus do not pay a fitness cost of *c*. In addition, in each interaction the microbes can be transmitted between interacting host, with probabilities *T*_*α*_, *T*_*β*_. Transmission and establishment of one microbe is independent of the other microbe, and when both occur, they occur simultaneously. The fitness of each host is determined according to the payoff matrix (Figure 1a in the main text). After all interactions take place, reproduction occurs. The next generation hosts are generated by considering the offspring as copies of the parent generation, and by choosing 10,000 hosts from the parent generation with a multinomial distribution, and with replacement (a parent can have more than one offspring). The probability to choose host *j* is its fitness divided by the sum of all the hosts’ fitness 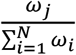.

#### Spatially-structured population

For the spatially-structured scenario we programmed an agent-based simulation where we consider a 2D 100 × 100 lattice, where each site is inhabited by one host. Each host is defined by the allele (*S*/*R*) and microbe (*α*/*β*) it carries. Cooperation and horizontal transmissions are defined similarly to the fully-mixed case, but differently from the fully-mixed model, the interactions are local. At each generation, each host (drawn in a random order) can interact with a randomly drawn neighbor host, where a neighbor is defined as any host from the 8 adjacent sites (or less if close to the edge of the lattice). We set the probability of each host to initiate an interaction at a given generation to be 1/2, so that the expected number of interactions per generation each host takes part in is approximately 1 (equal to 1 away from the borders). After all interactions take place, reproduction occurs. A new lattice grid of the same size is formed. Every site in the new lattice is inhabited by a replicate of a host from the neighborhood of that site in the original lattice, chosen randomly with probabilities proportional to the hosts’ fitness.

#### Mutations

We investigate the effects of mutations both in fully-mixed and spatially-structured populations. The mutations where modeled as a change in the allele or microbial population of an offspring host, relative to its parent. In the main text we show results of simulations with mutation rates of *µ* = 10^−4^ in all directions, namely equal probabilities for *α* → *β, β* → *α, S* → *R, R* → *S* mutations. In the stochastic simulations we examine populations of size *N* = 10,000, hence *µN* = 1. We also examined mutation rates of 10^−3^ and 10^−5^ and found qualitatively similar results (see Figure S3).

#### Stopping criteria of the simulation

When simulating populations without mutations, we let the simulation run until either one host type reaches fixation, or 5,000 generations. For simulations with mutations, we simply let the simulation run for 5,000 generations. We also examined the effect of prolonging the simulation time to 10,000 generations, and found very similar results, except in the fully-mixed population without mutations, where the range of parameters allowing the maintenance of cooperation narrowed, as expected (see Figure S4).

## Supplemental Material

**Supplemental Information** includes the derivations of the deterministic model, four supplemental figures, and one video.

**Supplementary video** shows the coevolutionary dynamic of a spatially-structured population along the generations, in one simulation run. On the left, the change in population composition is shown on the 2D-lattice, where each pixel represents one host. On the right, the change in host type frequencies in the same simulation run. Simulation parameters are: *T*_*β*_ = 0.25, *T*_*α*_ = 0.9 · *T*_*β*_, *c* = 0.05, *d* = 0.025, *b*/*c* = 8, no mutations.

## Authors’ contributions

O.L.-E. and L.H. designed the study and formulated the model. O.L.-E. derived the analytical equations and wrote the simulation code. O.L.-E. and L.H. analyzed the results and wrote the manuscript.

## Acknowledgements

We wish to thank Ranit Aharonov for comments on the manuscript.

