## Supplementary information for "Host-microbiome coevolution promotes cooperation in a rock-paper-scissor dynamic"

#### Supplementary Note 1 The host-microbe co-evolution model

Following are the equations that describe the population dynamics, as presented in the Methods section in the main text. We mark by $x_{\alpha S}, x_{\alpha R}, x_{\beta S}, x_{\beta R}$ the proportions of the host types in the current generation, and calculate $x'_{\alpha S}, x'_{\alpha R}, x'_{\beta S}, x'_{\beta R}$, the proportions of the host types in the next generation:

1. $$x'_{\alpha S} = f_1(x_{\alpha S}, x_{\alpha R}, x_{\beta S}, x_{\beta R}, c, b, \delta_\alpha, \delta_\beta, T_\alpha, T_\beta) = \frac{1}{\bar{\omega}}\big(x_{\alpha S}^2(1+b-c) + x_{\alpha S}x_{\alpha R}(1-c) + x_{\alpha S}x_{\beta S}\big[(1-T_\beta)(1-c) + T_\alpha(1+b)\big] + x_{\alpha S}x_{\beta R}(1-T_\beta)(1-c) + x_{\alpha R}x_{\beta S}T_\alpha\big)$$

2. $$x'_{\alpha R} = f_2(x_{\alpha S}, x_{\alpha R}, x_{\beta S}, x_{\beta R}, c, b, \delta_\alpha, \delta_\beta, T_\alpha, T_\beta) = \frac{1}{\bar{\omega}}\Big(x_{\alpha R}^2(1-\delta_\alpha) + x_{\alpha R}x_{\alpha S}(1+b-\delta_\alpha) + x_{\alpha R}x_{\beta S}(1-T_\beta)(1-\delta_\alpha) + x_{\alpha R}x_{\beta R}\,\big[(1-T_\beta)(1-\delta_\alpha) + T_\alpha(1-\delta_\beta)\big] + x_{\alpha S}x_{\beta R}T_\alpha(1+b-\delta_\beta)\Big)$$

3. $$x'_{\beta S} = f_3(x_{\alpha S}, x_{\alpha R}, x_{\beta S}, x_{\beta R}, c, b, \delta_\alpha, \delta_\beta, T_\alpha, T_\beta) = \frac{1}{\bar{\omega}}\Big(x_{\beta S}^2 + x_{\beta S}x_{\alpha S}\big[(1-T_\alpha)(1+b) + T_\beta(1-c)\big] + x_{\beta S}x_{\alpha R}(1-T_\alpha) + x_{\beta S}x_{\beta R} + x_{\alpha S}x_{\beta R}T_\beta(1-c)\Big)$$

4. $$x'_{\beta R} = f_4(x_{\alpha S}, x_{\alpha R}, x_{\beta S}, x_{\beta R}, c, b, \delta_\alpha, \delta_\beta, T_\alpha, T_\beta) = \frac{1}{\bar{\omega}}\Big(x_{\beta R}^2(1-\delta_\beta) + x_{\beta R}x_{\alpha S}(1-T_\alpha)(1+b-\delta_\beta) + x_{\beta R}x_{\beta S}(1-\delta_\beta) + x_{\beta R}x_{\alpha R}\big[T_\beta(1-\delta_\alpha) + (1-T_\alpha)(1-\delta_\beta)\big] + x_{\alpha R}x_{\beta S}T_\beta(1-\delta_\alpha)\Big)$$

where

5. $$\bar{\omega} = 1 - x_{\alpha R}\delta_\alpha - x_{\beta R}\delta_\beta + x_{\alpha S}(b-c)$$

#### Supplementary Note 2 Invasion Analysis

In order to analyze the stability of the four trivial equilibriums, where one type is at fixation and the others are extinct, we perform invasion analysis. We calculate the Jacobian of the system $(f_1, f_2, f_3, f_4)$, by deriving the functions with respect to the variables $x_{\alpha S}, x_{\alpha R}, x_{\beta S}, x_{\beta R}$. We then calculate the eigenvalues of the Jacobian in the four trivial equilibrium points.

We find that the Jacobian matrix at the equilibrium $x_{\alpha S} = 1$ is:

6. $$J|_{x_{\alpha S}=1} = \begin{pmatrix} \frac{2+b-c}{1+b-c} & \frac{1+\delta_\alpha - c}{1+b-c} & \frac{(1-T_\beta)(1-c)+T_\alpha(1+b)}{1+b-c} & \frac{(1-T_\beta)(1-c)+\delta_\beta}{1+b-c} \\ 0 & \frac{1+b-\delta_\alpha}{1+b-c} & 0 & \frac{T_\alpha(1+b-\delta_\beta)}{1+b-c} \\ 0 & 0 & \frac{(1-T_\alpha)(1+b)+T_\beta(1-c)}{1+b-c} & \frac{T_\beta(1-c)}{1+b-c} \\ 0 & 0 & 0 & \frac{(1-T_\alpha)(1+b-\delta_\beta)}{1+b-c} \end{pmatrix}$$

And the eigen values for invasion are $\frac{\mathbf{1+b-\delta_\alpha}}{\mathbf{1+b-c}}$, $\frac{\mathbf{(1-T_\alpha)(1+b)+T_\beta(1-c)}}{\mathbf{1+b-c}}$ and $\frac{\mathbf{(1-T_\alpha)(1+b-\delta_\beta)}}{\mathbf{1+b-c}}$.

By changing the order of the functions and the order of the derivations to $f_2, f_1, f_4, f_3$ and $x_{\alpha R}, x_{\alpha S}, x_{\beta R}, x_{\beta S}$ respectively, we find that the Jacobian matrix at the equilibrium $x_{\alpha R} = 1$ is:

7. $$J|_{x_{\alpha R}=1} = \begin{pmatrix} \frac{2-\delta_\alpha}{1-\delta_\alpha} & \frac{1+c-\delta_\alpha}{1-\delta_\alpha} & \frac{T_\alpha(1-\delta_\beta)+(1-T_\beta)(1-\delta_\alpha)+\delta_\beta}{1-\delta_\alpha} & 1-T_\beta \\ 0 & \frac{1-c}{1-\delta_\alpha} & 0 & \frac{T_\alpha}{1-\delta_\alpha} \\ 0 & 0 & \frac{T_\beta(1-\delta_\alpha)+\ (1-T_\alpha)(1-\delta_\beta)}{1-\delta_\alpha} & T_\beta \\ 0 & 0 & 0 & \frac{1-T_\alpha}{1-\delta_\alpha} \end{pmatrix}$$

And the eigen values for invasion are $\frac{\mathbf{1-c}}{\mathbf{1-\delta_\alpha}}$, $\frac{\mathbf{T_\beta(1-\delta_\alpha)+\ (1-T_\alpha)(1-\delta_\beta)}}{\mathbf{1-\delta_\alpha}}$ and $\frac{\mathbf{1-T_\alpha}}{\mathbf{1-\delta_\alpha}}$ .

By changing the order of the functions and the order of the derivations to $f_3, f_4, f_1, f_2$ and $x_{\beta S}, x_{\beta R}, x_{\alpha S}, x_{\alpha R}$ respectively, we find that the Jacobian matrix at the equilibrium $x_{\beta S} = 1$ is:

8. $$J|_{x_{\beta S}=1} = \begin{pmatrix} 2 & 1+\delta_\beta & 1+c+T_\beta(1-c)-T_\alpha(1+b) & 1+\delta_\alpha - T_\alpha \\ 0 & 1-\delta_\beta & 0 & T_\beta(1-\delta_\alpha) \\ 0 & 0 & T_\alpha(1+b)+(1-T_\beta)(1-c) & T_\alpha \\ 0 & 0 & 0 & (1-T_\beta)(1-\delta_\alpha) \end{pmatrix}$$

and the eigen values for invasion are: $\mathbf{1-\boldsymbol{\delta}_{\boldsymbol{\beta}}}$ , $\boldsymbol{T_\alpha(1+b)+(1-T_\beta)(1-c)}$ and $\boldsymbol{(1-T_\beta)(1-\delta_\alpha)}$ .

By changing the order of the functions and the order of the derivations to $f_4, f_3, f_2, f_1$ and $x_{\beta R}, x_{\beta S}, x_{\alpha R}, x_{\alpha S}$ respectively, we find that the Jacobian matrix at the equilibrium $x_{\beta R} = 1$ is:

9. $$J|_{x_{\beta R}=1} = \begin{pmatrix} \frac{2-\delta_\beta}{1-\delta_\beta} & 1 & \frac{T_\beta(1-\delta_\alpha)+(1-T_\alpha)(1-\delta_\beta)+\delta_\alpha}{1-\delta_\beta} & \frac{c-b+(1-T_\alpha)(1+b-\delta_\beta)}{1-\delta_\beta} \\ 0 & \frac{1}{1-\delta_\beta} & 0 & \frac{T_\beta(1-c)}{1-\delta_\beta} \\ 0 & 0 & \frac{T_\alpha(1-\delta_\beta)+(1-T_\beta)(1-\delta_\alpha)}{1-\delta_\beta} & \frac{T_\alpha(1+b-\delta_\beta)}{1-\delta_\beta} \\ 0 & 0 & 0 & \frac{(1-T_\beta)(1-c)}{1-\delta_\beta} \end{pmatrix}$$

and the eigen values are: $\boldsymbol{\frac{1}{1-\delta_\beta}}$ , $\boldsymbol{\frac{T_\alpha(1-\delta_\beta)+(1-T_\beta)(1-\delta_\alpha)}{1-\delta_\beta}}$ and $\boldsymbol{\frac{(1-T_\beta)(1-c)}{1-\delta_\beta}}$ .

To conclude, among all the invasion conditions, we find only one subset of conditions that their intersection maintains $0 < \delta_\beta, \delta_\alpha < c$, and allow a rock-paper-scissor cycle containing at least three host types, where $\alpha S$ is one of them. This result yields a dynamic where each of the four host types can be invaded by other host type, and thus all four trivial equilibria are not stable. The conditions are:

$$10a.\ \frac{b}{c} > \frac{1-T_\beta}{T_\alpha} + \frac{T_\beta - T_\alpha}{T_\alpha c}$$ (same as (1) in the main text)

$$10b.\ \delta_\alpha < c$$

$$10c.\ \delta_\beta > 0$$

$$10d.\ T_\alpha < \max\left(\delta_\alpha, T_\beta + \frac{\delta_\alpha - \delta_\beta}{1-\delta_\beta}(1-T_\beta)\right)$$

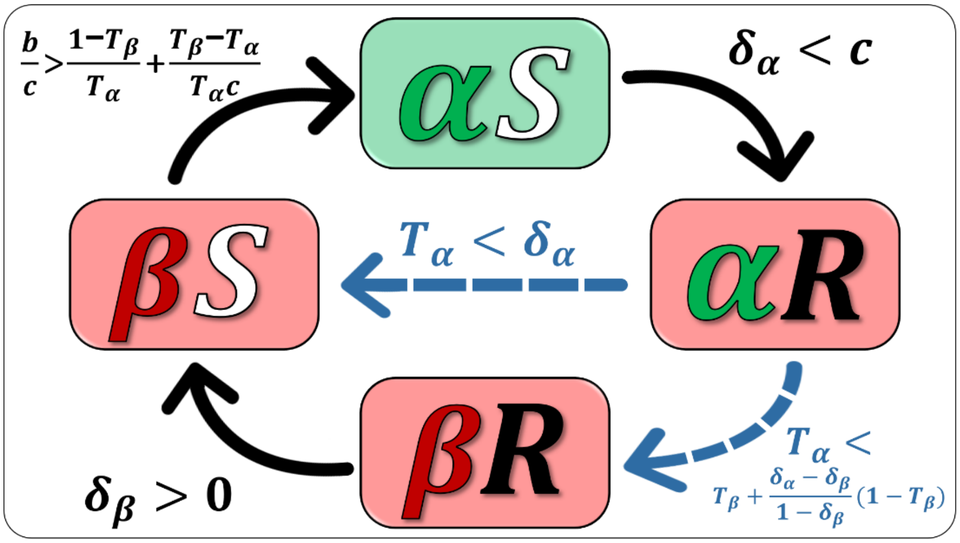


**Figure S1. Invasion dynamics based on the stability analysis performed in this section.** Illustrated are the set of conditions that maintain $0 < \delta_\beta, \delta_\alpha < c$, and allow a rock-paper-scissor dynamic. Note that when setting $\delta_\alpha = \delta_\beta = \delta$ we get the same dynamic as illustrated in figure 2c in the main text. We also note that $T_\beta + \frac{\delta_\alpha - \delta_\beta}{1-\delta_\beta}(1 - T_\beta) > T_\beta$ for $\delta_\alpha > \delta_\beta$, and thus the condition for $\alpha R \to \beta R$ is even milder than the one presented in the main text for $\delta_\alpha = \delta_\beta = \delta$.

When (10a-d) apply, there is no polymorphic equilibrium that involves two host types that share the same allele or microbe type. In addition, no polymorphic equilibrium that involves exactly two host types that carry different alleles and microbes can exist since horizontal transmission of the microbes will lead to generation of the other two host types as well. Similarly with three host types – no stable equilibrium can involve only three host types, since horizontal transmission will lead to the generation of the fourth host type. We thus conclude that when (10a-d) are maintained, there is no stable polymorphism on the boundaries of the 4-D standard simplex, hence the system must maintain all four host types. We find that when (10a-d) are maintained indeed the population either reach a stable polymorphic equilibrium, or oscillate chaotically.

### Supplementary Note 3 Equilibrium analysis

#### 3.1 General case of $\delta_\alpha \neq \delta_\beta$

We study the non-trivial equilibria of the system, where $x_i' = x_i$ for all $i \in \{\alpha S, \alpha R, \beta S, \beta R\}$. We begin by analyzing the equilibrium with respect to the alleles, where $x_{\alpha S}' + x_{\beta S}' = x_{\alpha S} + x_{\beta S}$ (based on equations 1 and 3). We denote $x_S = x_{\alpha S} + x_{\beta S}$, $x_R = x_{\alpha R} + x_{\beta R}$ and find that the proportion of allele $S$ can increase from one generation to the next only when:

$$11.\ \frac{x_{\alpha S}}{x_S} < \frac{\delta_\beta}{c} + \frac{\delta_\alpha - \delta_\beta}{c} \cdot \frac{x_{\alpha R}}{x_R}$$

and that a non-trivial equilibrium must satisfy:

$$12.\ \frac{x_{\alpha S}}{x_S} = \frac{\delta_\beta}{c} + \frac{\delta_\alpha - \delta_\beta}{c} \cdot \frac{x_{\alpha R}}{x_R}$$

Combining (12) and (1-5), and using a numeric solver we were able to find the polymorphic equilibrium (when such existed). The analysis showed that cooperation can evolve and be maintained even when $\delta_\beta < \delta_\alpha < c$.

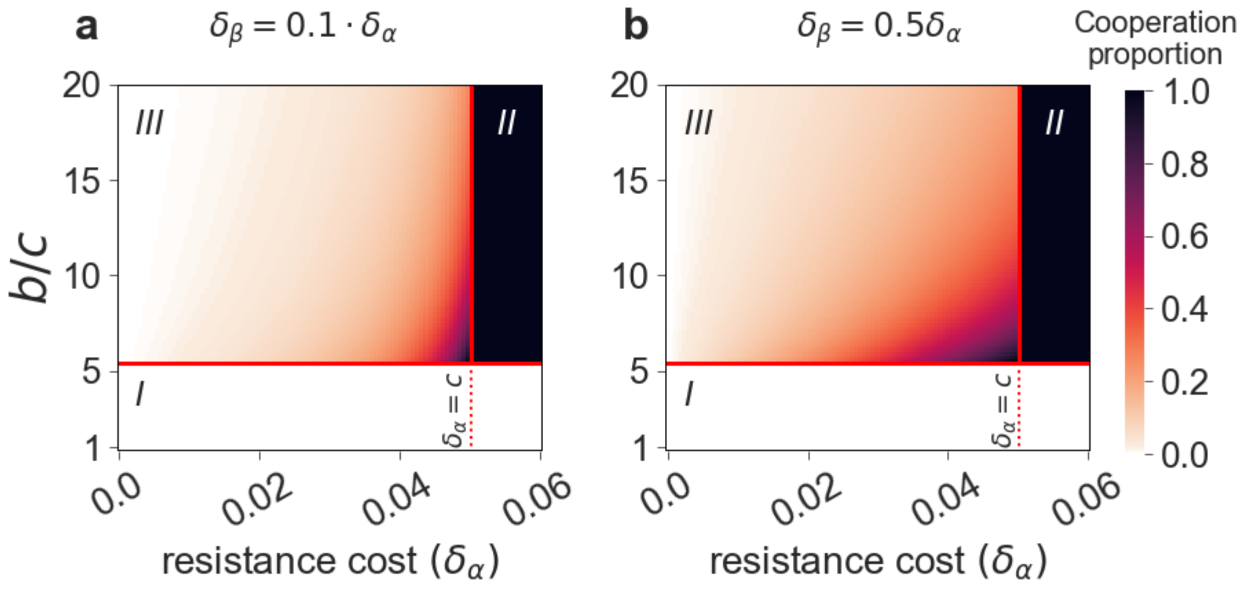

**Figure S2. Cooperation can be maintained also when $\delta_\beta < \delta_\alpha$.** We plot (color coded) the expected proportion of cooperative hosts ($\alpha S$) at equilibrium, as a function of $b/c$ (y-axis) and of $\delta_\alpha$ (x-axis) for $\delta_\beta = 0.1\delta_\alpha$ **(a)** and $\delta_\beta = 0.5\delta_\alpha$ **(b)**. Cooperation goes extinct when below the horizontal dashed line representing condition ***(1)*** of the main text (area ***I***, white). Above that threshold, cooperation can either go to fixation (when $\delta_\alpha > c$, area ***II***, black), or be maintained at intermediate levels (when $\delta_\alpha < c$, area ***III***). In the latter case, the proportion of cooperators increases with $\delta_\alpha$. We see that cooperation can be maintained even when $\delta_\beta < \delta_\alpha$, although the proportion of cooperators is lower for smaller values of $\delta_\beta/\delta_\alpha$ (compare Figure 2a from the main text, where $\delta_\alpha = \delta_\beta$ to this figure). $c = 0.05, T_\beta = 0.25, T_\alpha = 0.9 \cdot T_\beta$

##### 3.2 Special case of $\delta_\alpha = \delta_\beta = \delta$

We focused on the case where the cost of resistance is independent of the microbe the host carries, namely $\delta_\alpha = \delta_\beta = \delta$. In this case equation (12) becomes:

$$13.\quad \frac{x_{\alpha S}}{x_S} = \frac{\delta}{c}$$

A few insights arise from this equation. First, we can see that if $\delta > c$, the equilibrium cannot exist, since $x_{\alpha S} \leq x_S$ by definition. Second, this equation reveals that the proportion of cooperators at the equilibrium, is bounded by $\delta/c$. We further find that the proportion of allele $S$ can increase from one generation to the next only when (same as condition (2) in the main text):

$$14.\quad \frac{x_{\alpha S}}{x_S} < \frac{\delta}{c}$$

This means that when $\delta > c$, condition (14) applies for all $x_S > 0$ and therefore the proportion of allele $S$ will increase from one generation to the next until it reaches fixation. After $S$ reaches fixation, the dynamic is determined according to *(1)* of the main text: if the condition is maintained $\alpha S$ hosts will fixate, and otherwise, $\beta S$ hosts will fixate.

Using (13) and (1-5) we were able to find analytically two non-trivial equilibria of the system, and derive the exact solutions. We do not present here the exact expressions, as they are too long. Based on this analysis we generated figure 2 in the main text.

We continued by analyzing the validity of the non-trivial equilibria, namely that the proportions of each the four host types at equilibrium is positive and smaller than 1. We first note that when $\delta_\alpha = \delta_\beta = \delta$, conditions (10a-d) are simplified:

$15a.\ \frac{b}{c} > \frac{1-T_\beta}{T_\alpha} + \frac{T_\beta - T_\alpha}{T_\alpha c}$ (same as (1) in the main text)

$15b.\ \delta < c$

$15c.\ \delta > 0$

$15d.\ T_\alpha < \max(\delta, T_\beta)$

For any parameter set that maintain (15a-d) and that we've investigated, including the ones presented in figure 2 in the main text, only one polymorphic equilibrium was found. Screening more than $10^9$ parameter sets confirmed this finding. Since when (15a-d) are maintained the boundaries of the simplex are not stable (as explained in SI2), the system can either converge to the polymorphic equilibrium (as shown in Figures 3a,b in the main text), or oscillate chaotically around the equilibrium (as shown in Figures 3c,d in the main text). In any case, cooperation can evolve and be maintained at intermediate level.

We also found that there are some parameter sets that do not maintain (15a) or (15d), but still allow the existence of polymorphic equilibrium. Although we note, that the stability analysis shows that if one of the conditions (15a-d) is not maintained, then at least one of the trivial equilibria, is stable. In this case polymorphism cannot be globally stable, and if the system gets close enough to a stable trivial equilibrium it will get attracted and move towards it and remain there. When (15a-c) are maintained, but (15d) isn't (thus $T_\alpha > \max(\delta, T_\beta)$), there are some parameter sets for which equilibrium exists, but the dynamic depends on the initial proportions of the different host types. In this regime fixation of $\alpha R$ hosts is a stable equilibrium, and thus even though polymorphism can exist, it cannot be globally stable. Namely, some initial conditions of the population composition lead to polymorphism, while others lead to the fixation of $\alpha R$ hosts. When (15a) is not maintained we find that there are two types of dynamics. If (15d) is maintained, then fixation of $\beta S$ host is a stable equilibrium and it is the only one among the trivial equilibria. In this case the system drives the population towards fixation of $\beta S$ hosts (verified numerically over more than $10^9$ parameter sets). If on the other hand both (15a) and (15d) are not maintained, there are two stable trivial equilibria in the system – the fixation of $\beta S$ hosts and the fixation of $\alpha R$ hosts. In this range of parameters there is also a polymorphic equilibrium. Nevertheless, in any parameter set that we've investigated, stability analysis revealed that the polymorphic equilibrium is unstable.

#### Supplementary Note 4 Stochastic Simulations

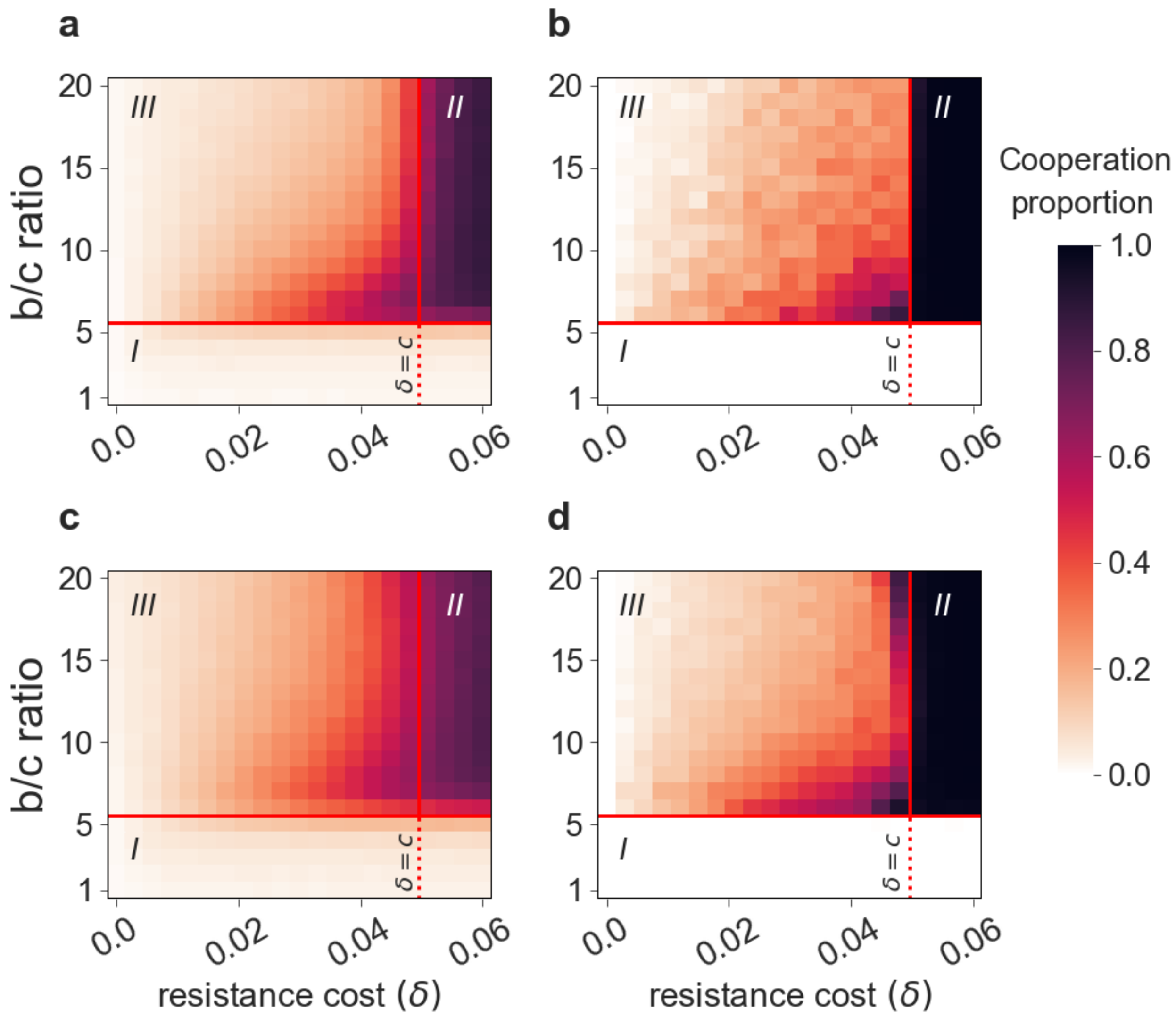


**Figure S3. Cooperation is maintained under varied mutation rates.** Similarly to figure 4 in the main text, the proportion of cooperators after 5,000 generations is plotted as a function of the $b/c$ ratio on the y-axis and $\delta$ on the x-axis. The color of each site represents the average of 100 stochastic simulation runs. Panels **(a)** and **(b)** shows the results of fully-mixed populations, while panels **(c)** and **(d)** show results of spatially-structured populations. For panels **(a)** and **(c)** we set mutation rate $\mu = 10^{-3}$ in all directions ($\alpha \leftrightarrow \beta$ $and$ $S \leftrightarrow R$), while for panels **(b)** and **(d)** we used $\mu = 10^{-5}$ in all directions. Note that in panels **(a)** and **(c)**, when $\mu = 10^{-3}$, limited cooperation is maintained even below the $b/c$ threshold derived from condition (1) of the main text, due to mutation-selection balance. Simulation parameters: $T_\beta = 0.25, T_\alpha = 0.9T_\beta, c = 0.05$

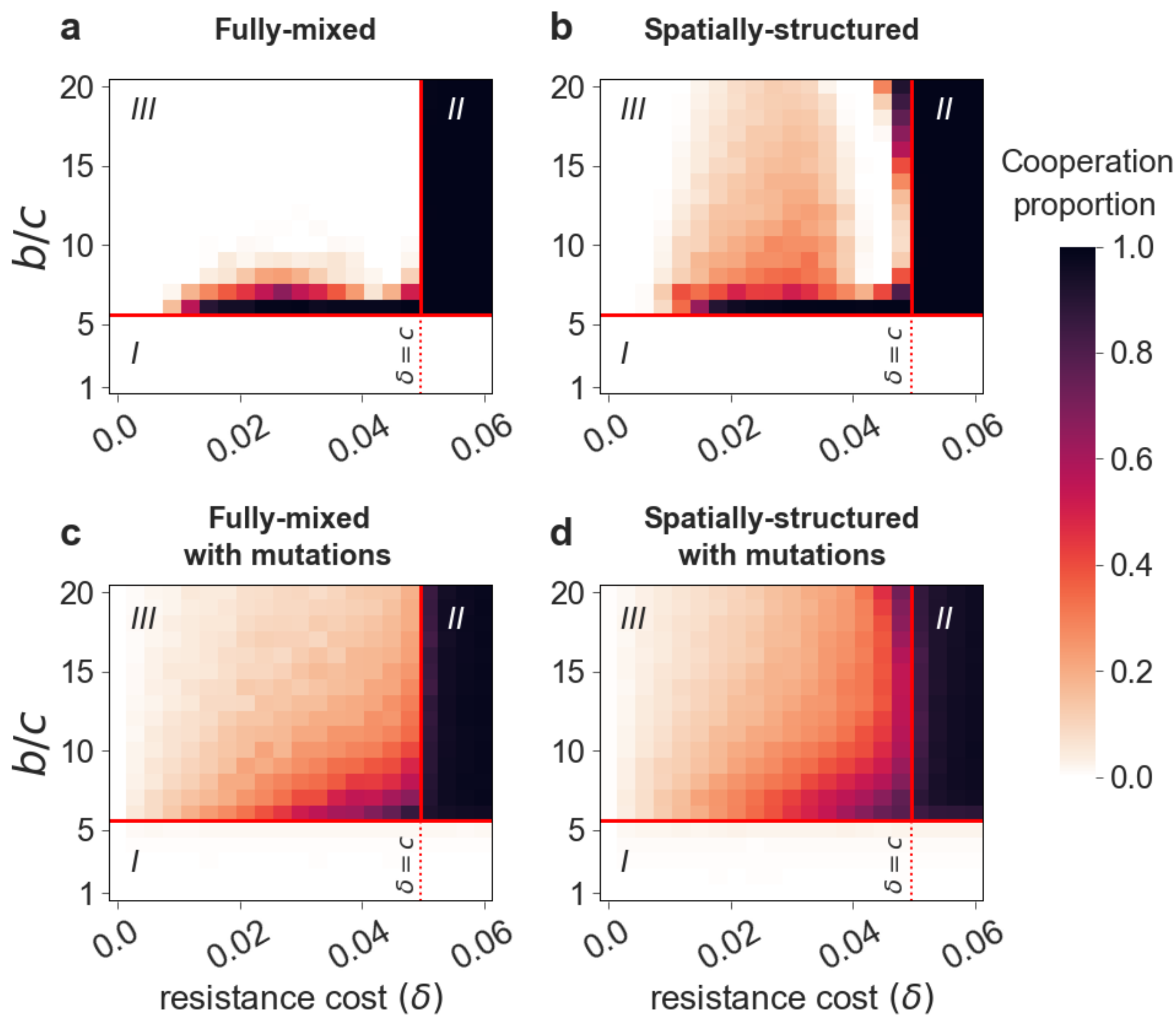


**Figure S4. Mutations and spatial structure help maintaining cooperation in the face of host resistance and finite populations.** This figure is similar to Figure 4 in the main text except for the stopping condition. In this figure we show the proportion of cooperators after 10,000 generations. The color of each site represents the average of 100 stochastic simulation runs. It can be seen that the results are very similar to the ones presented in Figure 4 in the main text, except for panel **(a)**, where the range of parameters allowing the maintenance of cooperation is narrower here.
